# Repository-scale Co- and Re-analysis of Tandem Mass Spectrometry Data

**DOI:** 10.1101/750471

**Authors:** Alan K. Jarmusch, Mingxun Wang, Christine M. Aceves, Rohit S. Advani, Shaden Aguire, Alexander A. Aksenov, Gajender Aleti, Allegra T. Aron, Anelize Bauermeister, Sanjana Bolleddu, Amina Bouslimani, Andres Mauricio Caraballo Rodriguez, Rama Chaar, Roxana Coras, Emmanuel O. Elijah, Madeleine Ernst, Julia M. Gauglitz, Emily C. Gentry, Makhai Husband, Scott A. Jarmusch, Kenneth L. Jones, Zdenek Kamenik, Audrey Le Gouellec, Aileen Lu, Laura-Isobel McCall, Kerry L. McPhail, Michael J. Meehan, Alexey V. Melnik, Riya C. Menezes, Yessica Alejandra Montoya Giraldo, Ngoc Hung Nguyen, Louis Felix Nothias, Mélissa Nothias-Esposito, Morgan Panitchpakdi, Daniel Petras, Robert Quinn, Nicole Sikora, Justin J.J. van der Hooft, Fernando Vargas, Alison Vrbanac, Kelly Weldon, Rob Knight, Nuno Bandeira, Pieter C. Dorrestein

## Abstract

Metabolomics data are difficult to find and reuse, even in public repositories. We, therefore, developed the Reanalysis of Data User (ReDU) interface (https://redu.ucsd.edu/), a community- and data-driven approach that solves this problem at the repository scale. ReDU enables public data discovery and co- or re-analysis via uniformly formatted, publicly available MS/MS data and metadata in the Global Natural Product Social Molecular Networking Platform (GNPS), consistent with findable, accessible, interoperable, and reusable (FAIR) principles.^1^

## Results and Discussion

Many simple but important questions can be asked using repository-scale public data. For example, what human biospecimen or sampling location is best for detecting a given drug? Or what molecules are found in humans <2 years old? Current metabolomics repositories typically require manual navigation and conversion of thousands of different vendor-formatted files with inconsistent metadata formats, and developing data integration algorithms, greatly complicating analyses.

ReDU addresses FAIR principles by enabling users to find and choose files (Fig 1a). This is possible because ReDU formats sample information consistently via a template and drag-and-drop validator backed by standard controlled vocabularies and ontologies (*e.g.* NCBI taxonomy,^2^ UBERON^3^, Disease Ontology^4^ and MS ontology), and includes geographical location (important for natural products and environmental samples). ReDU automatically uses all public data in the GNPS/MassIVE repository that has the corresponding ReDU-compliant sample information. 34,087 files in GNPS are ReDU-compatible including natural and human-built environments, human and animal tissues, biofluids, food, and other data from around the world (Fig 1f), analyzed using different instruments, ionization methods, sample preparation methods, etc. From the 103,230,404 million MS/MS spectra included in ReDU, 4,528,624 spectra were annotated (rate of 4.39% with settings yielding ~1% FDR) as one of 13,217 unique chemicals (**Table S1**).^5,6,7^

**Fig 1.**
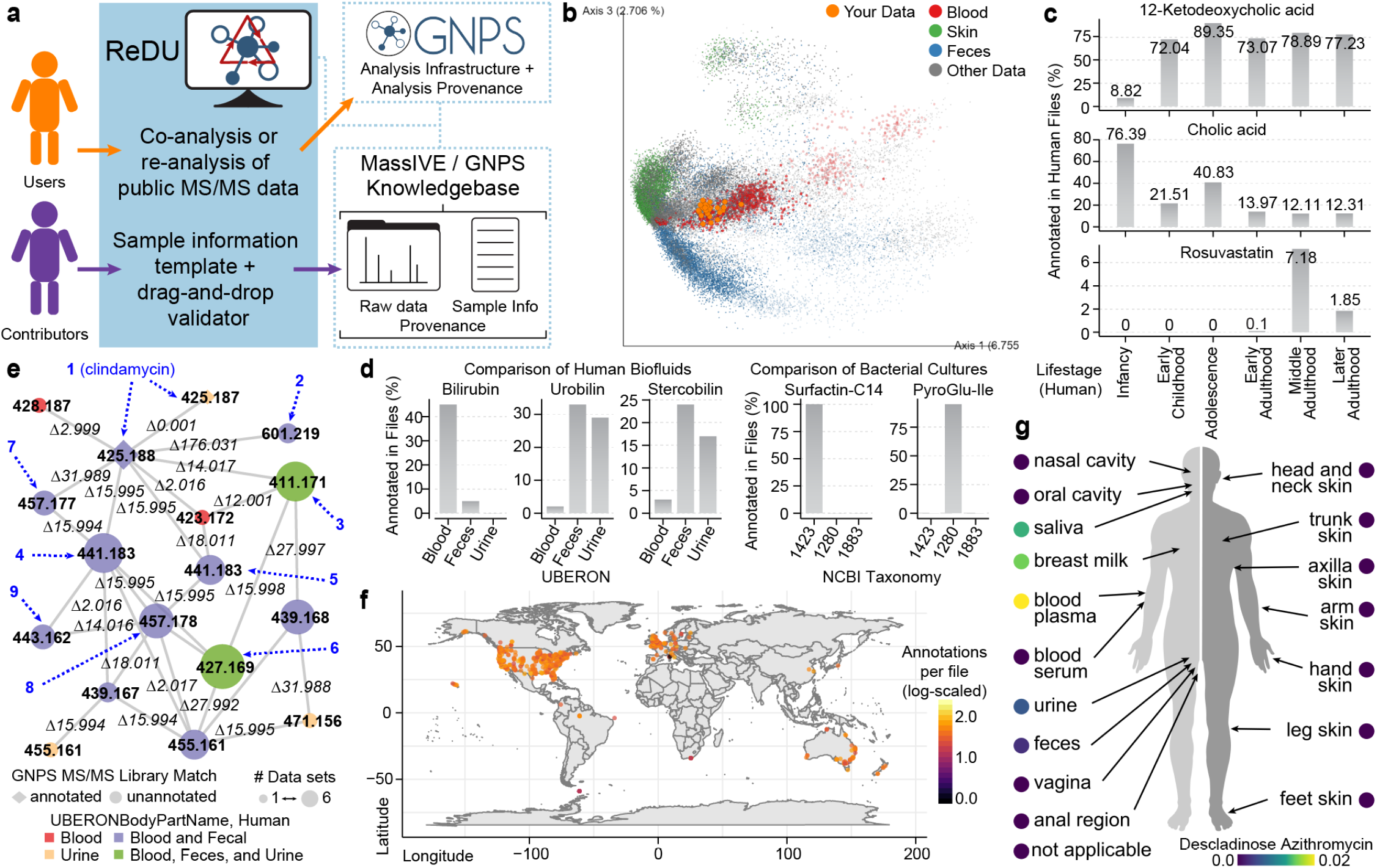
ReDU workflow and illustrative applications. **(a)** The ReDU framework. **(b)** Users can co-analyze their data via projection onto public data visualized using EMPeror^11^. PCA was performed on GNPS annotations (level 2/3).^7^ Human blood plasma samples (orange) from rheumatoid arthritis patients. Sample points size and color were set using UBERON ontology and opacity was set using NCBI taxonomy filtering (your data, 1.0; 9606|Homo sapiens, 0.7; and all other data, 0.25). **(c)** Metadata filters were used to select human fecal samples (NCBI taxonomy, 9606|Homo sapiens; UBERON, feces) and launch sample information enrichment. Chemical explorer was used to select 12-Ketodeoxycholic acid, cholic acid, and rosuvastatin. Lifestages: Infancy (<2 yrs), n=1859; Early Childhood (2 yrs < x ≤ 8 yrs), n=93; Adolescence (8 yrs < x ≤ 18 yrs), n=169; Early Adulthood (18 yrs < x ≤ 45 yrs), n=995; Middle Adulthood (45 yrs < x ≤ 65 yrs), n=933; and Later Adulthood (> 65 yrs), n=325. **(d)** Metadata filters were used to select human blood, feces, and urine into different groups and launch chemical enrichment analysis. Bilirubin, urobilin, and stercobilin are illustrative of the chemical differences between the groups. Similarly, bacterial cultures of 1423|Bacillus subtilis (n=89), 1280|Staphylococcus aureus (n=49), and 1883|Streptomyces (n=7) were selected into groups using filters. Surfactin-C14 and PyroGlu-Ile were two exemplar chemicals observed to be different between groups. **(e)** Human blood, feces, and urine were selected using ReDU metadata filters and re-analyzed together using molecular networking. A portion of the molecular network associated with clindamycin is displayed. Nodes are colored by the UBERON ontology in which it was detected: node size reflects the number of MassIVE datasets in which it was detected, node shape indicates whether annotated via library search (annotated, diamond or unannotated, circle). Putatively annotated clindamycin metabolites are indicated using dashed arrows and numbers, blue, corresponding to the proposed structures (**1-9**). **(f)** ReDU sample locations (includes public data from samples of environmental, natural products and other cohorts for which this information is provided) colored by number of annotations per file (latitude and longitude), log_10_-scale, using the ReDU database. **(g)** ReDU database (filtered to include human data) analyzed to visualize the distribution of chemical annotations tagged as drug or drug metabolite using ‘ili.^12^ Descladinose azithromycin was detected in blood plasma, breast milk, saliva, urine, and fecal samples. The number of annotations are divided by the number of files per sample type (*i.e.* UBERON). A distribution map of all drugs in ReDU is provided.

Uniformity of data and sample information in ReDU enables metadata-based and repository-scale analyses (Fig. 1b–g). Chemical explorer enables selection of a molecule and retrieval of its associations with the metadata, *i.e.* sample information association. For instance, selecting 12-ketodeoxycholic acid (filtering to include human feces) revealed it was observed after infancy (Fig 1c), whereas cholic acid displayed the opposite trend, coupled to the developing microbiome. Similarly, rosuvastatin was found in adults matching prescription demographics. Another approach enabled is chemical enrichment analysis. For example, human blood, feces, and urine differed by bilirubin, urobilin, and stercobilin (Fig 1d). Bilirubin was more frequently annotated in blood, and urobilin and stercobilin were most often annotated in feces.^8^ Similarly, comparison of bacterial cultures revealed differences in annotation of surfactin-C14 (observed in *Bacillus subtilis*) and cholic acid (observed in *Streptomyces*). ReDU enables reanalysis based on metadata-selected files for molecular networking.^5,9,10^ Re-analyzing human blood plasma and serum, urine, and fecal samples, networked 5,053,666 MS/MS spectra (~5.6% annotated) and included annotations to clindamycin. Clindamycin’s (**1**) molecular family matched multiple datasets and sample types (Fig 1e). Using propagation through molecular networking (e.g. delta mass and MS/MS spectral interpretation), we annotated clindamycin metabolites (**2**-**9**) **Fig S1**-**S2, Table S2**-**S3**. *N*-desmethylclindamycin sulfoxide (**6**) was observed in multiple sample types across six different datasets. At the repository scale, we can map sample geographical location and identify the number of chemical annotations for each sample (Fig 1f), or locate specific molecules of interest (*e.g.* drugs) by mapping on the human body offline (Fig 1g, **Video S1**). These are representative analyses uniquely enabled by the ReDU infrastructure.

ReDU makes public MS/MS data FAIR and connects MassIVE (a data repository recommended by Nature publishing journal for metabolomics and proteomics data) to GNPS (an analysis environment), thereby integrating public data deposition, sample information curation, and data analysis. ReDU’s utility will continue to grow as more data is uploaded to GNPS/MassIVE,and public reference libraries expand, making ReDU a resource developed by the community and FAIR for the community.

## Author Contributions

AKJ, MW, and PD developed the ReDU concept.

AKJ, MW, and CMA wrote code and engineered the ReDU infrastructure.

AKJ, CMA, RSA, SA, AAA, GA, AA, AB, SB, AB, AMCR, RC, EOE, JJJvdH, JMG, ECG, MH, KJ, ZK, ALG, AL, LIM, KLM, MJM, AVM, RCM, YAM, NHN, LFN, ME, MNE, MP, DP, RQ, NS, FV, AV, and KW curated metadata enabling ReDU.

AKJ, MW, CMA, SAJ, LM, ME, JJJvdH, JMG, MP and PCD tested the ReDU infrastructure and provided feedback.

AKJ, MW, CMA, ME, JJJvdH, RK, NB, and PCD wrote and edited the manuscript.

RK, NB, and PCD provided supervision and funding support.

## Ethics Declaration

Pieter C. Dorrestein is a scientific advisor for Sirenas LLC.

Mingxun Wang is a consultant for Sirenas LLC and the founder of Ometa labs LLC.

Alexander Aksenov is a consultant for Ometa labs LLC.

## Acknowledgments

The authors would like to thank the individuals involved in the funding, administration, sample collection, and data acquisition of the public data used in ReDU. The authors recognize the financial support of the U.S. National Institutes of Health (P41 GM103484, R03 CA211211, and R01 GM107550), Sloan Foundation (RK), Gordon and Betty Moore Foundation (PD, NB, KLM), American Society for Mass Spectrometry (AKJ), NSF grant IOS-1656481 (PCD and AMCR), Netherlands eScience Center No. ASDI.2017.030 (JJJvdH), Krupp Endowed Fund (RC)the Department of the Navy, Office of Naval Research Multidisciplinary University Research Initiative (MURI) Award, Award number N00014-15-1-2809 and the University of California, San Diego Center for Microbiome Innovation SEED grants.

## Data Availability

All curated sample information can be downloaded from the ReDU homepage (https://redu.ucsd.edu/) by selecting “Download Database.” The current version of the ReDU information is archived in the GNPS/MassIVE (gnps.ucsd.edu) repository. The accession number is MSV000084206.

## Code Availability

All code for ReDU is available in GitHub (https://github.com/mwang87/ReDU-MS2-GNPS) with corresponding documentation (https://github.com/mwang87/ReDU-MS2-Documentation).

## Notes

https://redu.ucsd.edu/

https://gnps.ucsd.edu/

https://massive.ucsd.edu

