## Supplemental Information for "Repository-scale Co- and Re-analysis of Tandem Mass Spectrometry Data"

### Supplemental Materials

**Video S1.** Video depicting the distribution of drugs annotated in ReDU in human samples. The video can be accessed via the following link: https://www.youtube.com/watch?v=dzAqjBNmqPU&feature=youtu.be

### Supplemental Methods

#### ReDU Content

The homepage of ReDU (https://redu.ucsd.edu/) is the launch point for different analyses, centered around “Analyze Your Data” or “Analyze Public Data”. It also links to “Documentation”, “How to Contribute Data”, “ReDU Sample Information Validator”, “Download Database”, and “File Query - Sample Information”. The “Documentation” option (**Fig S3a**) links to the ReDU documentation, and the “How to Contribute Data” option (**Fig S3b**) links to the subsection of documentation which list the steps necessary to contribute data to ReDU. The “ReDU Sample Information Validator” (**Fig S3c**) links to a drag-and-drop validator (**Fig S4**) that verifies that the sample information template (**Fig S5**) required for data contribution adheres to the required formatting and terms in a controlled vocabulary (additional terms must be submitted via GitHub). “Download Database” (**Fig S3d**) downloads all the sample information included in ReDU in a tab-separated text file. “File Query - Sample Information” (**Fig S3e**) links to a page that displays all sample information associated with a filename in MassIVE. The ReDU server is built using the python flask framework, SQLite, and a Vue.js front end.

#### Data and Sample Information Contribution

Data files (.mzXML or .mzML) and a ReDU-validated sample information table are necessary for inclusion of data in ReDU and must be uploaded to a public MassIVE data set. A sample information template (**Fig S5**) and validator (**Fig S4**) are provided. Detailed step-by-step instructions can be found in the ReDU documentation (https://mwang87.github.io/ReDU-MS2-Documentation/HowtoContribute/).

#### Analyze Your Data: Co-analysis of Your Data to Metadata Filtered Public Data via Multivariate Analysis

Users can co-analyze their data onto an EMPeror plot of all data in ReDU (**Fig S3f**). Principal component analysis (PCA) was performed on the counts of each chemical annotation from GNPS spectral library matching using GNPS' default parameters (https://ccms-ucsd.github.io/GNPSDocumentation/librarysearch/). This search was performed on the MS/MS product ion scans *de novo* from files located in MassIVE. PCA was performed in python with scikit-learn. The eigenvector matrix was retained and used to calculate the location of the projected points. Projection was performed by multiplying the annotations for each file (vector) by the eigenvectors in order to calculate the location of data points in the precalculated coordinate frame. Users submit their data by providing a GNPS task ID into the field. GNPS library search, GNPS molecular networking, and GNPS feature-based molecular networking are compatible. It is encouraged that default library search parameters of GNPS-based networking jobs with minimum 6 ions required to match at cosine >0.7 be used. The task ID provides the information required to calculate the coordinates for the projection of samples onto the precalculated PCA plot (visualized using EMPeror) of all ReDU data. The EMPeror plot is interactive and the user has plotting options (including the axes and the color of points based on sample information), filtering options, and can rescale data. Furthermore, the user can highlight their data using the "Your Data" term in the "type" category; we suggest using this column to change the scale or opacity of the sample points to visualize your data. Clicking on any of the points in the plot results in the filename being displayed in the bottom-left corner. The entire PCA plot can be saved as an image file. Further download options will follow ReDU software releases.

##### Projection of Human Blood Samples Example - Fig 1b

Human blood plasma samples not yet entered in ReDU at the time of analysis were subjected to a GNPS library search using default parameters; the data and illustrative library search can be accessed using the following link (https://gnps.ucsd.edu/ProteoSAFe/status.jsp?task=f39c94cb7afe4568950bf61cdb8fee0d). The task ID was entered using the "Compare Your Data to Public Data via Multivariate Analysis" option (https://redu.ucsd.edu/comparemultivariate) resulting in the EMPeror plot displayed in Fig 1. The example button will populate the field with the task ID used to generate the figure. The following settings were used to create the image. Points were scaled using the UBERONBodyPartName category and globally scaled to 1.3 with the exception of blood, blood plasma, blood serum which were scaled to 2.5 and the projected data scaled to 5. The opacity was set to 0.25 globally using the NCBITaxonomy column, and the values for the projected data were set to 1 and all 9606|Homo sapiens data set to 0.7. Points were colored based on UBERONBodyPartName. All points were set to grey (#808080) except skin samples (green, #4daf4a), blood samples (red, #e41a1c), feces (blue, #377eb8), and the projected data (orange, #ff7f00). A .json file (settings file) has been provided to reproduce the plot by uploading it in the "load saved settings" option. This example is only intended to illustrate that blood samples cluster closely with other blood samples already in the ReDU database.

#### Analyze Your Data: Co-analyze Your Data with Metadata Filtered Public Data at GNPS With Molecular Networking

Users can co-analyze their data with public data by clicking the "Co-analyze Your Data with Public Data at GNPS" text, which links to the file selector (**Fig S3g** and **Fig S6**). The file selector allows one to select (and filter) files based on the sample information and place multiple types of files into one of 6 different groups (*i.e.* G1-G6) for molecular networking via GNPS. Library search without molecular networking, providing annotations only, can be formed via GNPS; however, all files should be placed in G1 as groups are not supported. Once the user has selected the public files they wish to include, a click of the "Co-analyze with GNPS Molecular Networking" or "Co-analyze with GNPS library search" button will load the public files into a GNPS molecular networking or GNPS library search launch page, respectively, at which point the user can add their own files to the appropriate group and submit the job. Details on molecular networking and library search can be found in the GNPS documentation (https://ccms-ucsd.github.io/GNPSDocumentation/). A maximum of 5000 files for molecular networking is suggested. If greater than 5000 files are to be co-networked then we suggest to contact the authors as more compute resources will need to be allocated.

#### Analyze Public Data: Explore Multivariate Analysis of Public Data

The PCA plot visualized using Emperor used in the "Compare Your Data to Public Data via Multivariate Analysis" section can be accessed (**Fig S3h**). Additional information for plotting and using Emperor can be found on GitHub (https://github.com/biocore/emperor).

#### Analyze Public Data: Explore Sample Information Association

Users can explore the association between chemical annotations and the sample information associated with the files in which they are detected (**Fig S3i**). All chemical annotations are tabulated and can be explored (**Fig S7**). A text search can be performed in the browser for a particular chemical of interest. The sample information association for a chemical can be accessed by clicking the "View Association" button as well as a list of files in which the chemical was found by clicking the "View Files" button. The sample information is tabulated for the selected chemical and ranked based on the proportion of files associated with a sample information term.

#### Analyze Public Data: Re-analyze Public Data at GNPS

##### Molecular Networking and Library Search

Users can re-analyze with public data by clicking the “Reanalyze Public Data at GNPS” text (**Fig S3j**), which links to the file selector, **Fig S6,** (shared by the “Co-analyze Your Data with Public Data at GNPS” section). The use of the file selector is detailed previously in the “Co-analyze Your Data with Public Data at GNPS” section. Upon completion of data selection, the user can launch the “Re-analyze with GNPS Molecular Networking” or “Re-analyze with GNPS library search” buttons which populate the GNPS molecular networking or GNPS library search launch page, respectively. The suggested parameters for molecular networking and library search are detailed in the GNPS documentation (https://ccms-ucsd.github.io/GNPSDocumentation/). A maximum of 5000 files for molecular networking is suggested.

**Molecular Networking Example - Fig 1e**

Molecular networking was performed in GNPS after selecting human blood plasma and serum (n=711), human urine (n=307), and human fecal (n=5114) files in the ReDU file selector (https://gnps.ucsd.edu/ProteoSAFe/status.jsp?task=a75aa494e927481dae6de12608e5e4a0). The data were filtered by removing all MS/MS peaks within ± *m/z* 17 of the precursor *m/z*. MS/MS spectra were window filtered by choosing only the top 6 peaks in the ± *m/z* 50 window throughout the spectrum. The data was then clustered with MS-Cluster with a precursor *m/z* tolerance of 0.02 and a MS/MS fragment ion (i.e. product ion) *m/z* tolerance of 0.02 to create consensus spectra. Further, consensus spectra that contained less than 5 spectra were discarded. A network was then created where edges were filtered to have a cosine score above 0.7 and more than 5 matched peaks. Further edges between two nodes were kept in the network if and only if each of the nodes appeared in each other's respective top 10 most similar nodes. The spectra in the network were then searched against GNPS' spectral libraries. The library spectra were filtered in the same manner as the input data. All matches kept between network spectra and library spectra were required to have a score above 0.7 and at least 5 matched peaks. The network was opened in Cytoscape (3.7.1), cytoscape.org,[1] and the networks were output as a .pdf and assembled in Adobe Illustrator. The molecular networking component associated with clindamycin was analyzed using the in-browser network visualization (https://gnps.ucsd.edu/ProteoSAFe/result.jsp?view=network_displayer&componentindex=2892&task=a75aa494e927481dae6de12608e5e4a0#%7B%7D) which was used to download the spectra displayed in **Fig S2**.

**Chemical Enrichment**

Users can compare the occurrence of chemical annotations between two or more groups populated in the file selector by clicking the “Launch Chemical Enrichment Analysis” button after data selection. GNPS chemical annotations are tabulated with the number of files in which they are found (and the percentage of files) in each group (G1-G6), **Fig S8**. The information displayed are precalculated (same information used for PCA) using default library search parameters.

**Chemical Enrichment Example - Fig 1d**

The file selector was used to filter only human files (NCBItaxonomy = 9606|Homo sapiens). Blood plasma and blood serum files were selected into G1, urine files were selected into G2, and fecal files were selected into G3. Chemical enrichment, a tabulation of chemicals and corresponding counts (*i.e.* number of times annotated) in each group, was launched. The resulting table displayed on the ReDU website was copied and pasted into Excel (Microsoft). The columns were arranged such that the number of files and the percentages for each group were split into different columns, and the percentages were converted into numeric proportions. The data file was saved as a tab delimited text file and imported into R for plotting.

The file selector was used to filter only bacterial cultures (SampleType = culture_bacterial). 1423|Bacillus subtilis (n=89) files were selected into G1, 1280|Staphylococcus aureus (n=49) files were selected into G2, and 1883|Streptomyces (n=7) files were selected into G3. The NCBITaxonomy metadata category was used for file selection. Chemical enrichment was launched. The resulting table displayed on the ReDU website was copied and pasted into Excel (Microsoft). The columns were arranged such that the number of files and the percentages for each group were split into different columns, and the percentages were converted into numeric proportions. The data file was saved as a tab delimited text file and imported into R for plotting.

**Sample Information Association**

The sample information association option is identical to that described in the “Analyze Public Data: Explore Sample Information Association” section, expect that only the files selected in the file selector and placed into group 1 (G1) are considered in the calculation.

**Sample Information Association Example - Fig 1c**

The file selector was used to filter only human files (NCBItaxonomy = 9606|Homo sapiens) and fecal samples were selected into G1 using the UBERONbodypartname. Sample information was launched. The resulting webpage was searched using Ctrl+F and searching for the chemicals to plot. The view associations button was clicked for each. The resulting table displayed on the ReDU website, similar to that displayed in **Fig S7**, was copied and pasted into Excel (Microsoft). All associations were tabulated in a single spreadsheet, and an additional column indicating the chemical was added. The data file was saved as a tab delimited text file and imported into R for plotting.

#### Illustrative Use of the ReDU Database: Map - Fig 1f

The ReDU information (MSV000084206) was downloaded and the latitudinal and longitudinal data were cleaned of any non-adherent formatting. The number of unique files associated with each latitude and longitude coordinate was calculated as well as the number of chemical annotations. The sum of the chemical annotations per latitude and longitude coordinate was divided by the number of unique files associated with the coordinates. Files lacking coordinates were removed. The values were $\log_{10}$ scaled to aid in visualization. The data were plotted in R (“ggmap” and “map” packages were used) onto a world map.

#### Illustrative Use of the ReDU Database: Drugs in Human Samples Visualized via ‘ili - Fig 1g

The ReDU information (MSV000084206) was downloaded and merged with the sample information database. A list of curated tags was generated from the curated source information table (provided). The files associated with humans were included and the chemical annotations associated with drugs or drug metabolites, putatively, were included. The number of chemical annotations per UBERON body part were divided by the number of files included for each body part. An image of an androgynous human was created in Illustrator (Adobe) and saved as a .png. The pixel coordinates associated with each label were tabulated by UBERON ontology name and merged with the ReDU drug table. The resulting file was exported as a .csv for use in ‘ili. These files, and a .json file that will reproduce the illustrative example in the manuscript, are available https://github.com/mwang87/ReDU-MS2-GNPS/tree/master/examples. The results were compiled into a video (https://www.youtube.com/watch?v=dzAqjBNmqPU&feature=youtu.be).

##### MS/MS Reference Library Hits

All annotation information was downloaded from ReDU and processed in R. Script is available on GitHub in the examples folder. The data were grouped by the MS/MS reference library available in GNPS (*e.g.* GNPS-LIBRARY), and spectral matches were tallied (excluding multiple hits to the same CCMSlib ID in the same file). The number of reference spectra were counted. (https://proteomics2.ucsd.edu/ProteoSAFe/result.jsp?task=ba6a5b6a1c0946b3a641c67ad59fb2df&view=production_library_sizes#%7B%22table_sort_history%22%3A%22main.number_spectra_dsc%22%7D). The information is tabulated in **Table S1**.

#### Supplemental Tables

**Table S1**. Tabulated MS/MS reference libraries used in GNPS, the number of spectra in each library, and the number of ReDU annotations per reference library.

| MS/MS Reference Library Name Available in GNPS | Number of Reference Spectra | Number of ReDU Spectral Matches |
|---|---|---|
| GNPS-NIST14-MATCHES | 5763 | 944273 |
| PNNL-LIPIDS-POSITIVE | 30582 | 298000 |
| MONA | 49241 | 278184 |
| GNPS-NIH-NATURALPRODUCTSLIBRARY_ROUND2_POSITIVE | 5796 | 194438 |
| GNPS-LIBRARY | 4697 | 150457 |
| MASSBANK | 11999 | 110935 |
| RESPECT | 7112 | 33684 |
| GNPS-EMBL-MCF | 585 | 31082 |
| BILELIB19 | 177 | 25392 |
| HMDB | 2235 | 16827 |
| GNPS-SELLECKCHEM-FDA-PART1 | 2388 | 12760 |
| MASSBANKEU | 1492 | 9239 |
| CASMI | 568 | 9164 |

| | | |
|---|---|---|
| GNPS-COLLECTIONS-MISC | 46 | 5757 |
| GNPS-SELLECKCHEM-FDA-PART2 | 656 | 5050 |
| DEREPLICATOR_IDENTIFIED_LIBRARY | 379 | 4840 |
| GNPS-NIH-SMALLMOLECULEPHARMACOLOGICALLYACTIVE | 1460 | 4625 |
| GNPS-NIH-NATURALPRODUCTSLIBRARY | 1267 | 4474 |
| GNPS-NIH-CLINICALCOLLECTION1 | 377 | 3512 |
| PNNL-LIPIDS-NEGATIVE | 16142 | 3126 |
| SUMNER/Bruker | 904 | 2742 |
| GNPS-COLLECTIONS-PESTICIDES-POSITIVE | 653 | 2707 |
| LDB_NEGATIVE | 226 | 2632 |
| GNPS-NIH-NATURALPRODUCTSLIBRARY_ROUND2_NEGATIVE | 1863 | 1667 |
| GNPS-FAULKNERLEGACY | 127 | 1322 |
| LDB_POSITIVE | 83 | 745 |
| GNPS-PRESTWICKPHYTOCHEM | 143 | 689 |
| GNPS-NIH-CLINICALCOLLECTION2 | 195 | 671 |
| MMV_POSITIVE | 110 | 585 |
| MIADB | 172 | 571 |
| GNPS-COLLECTIONS-PESTICIDES-NEGATIVE | 76 | 9 |

**Table S2.** Tabulated information for putatively annotated clindamycin-related chemicals in the clindamycin molecular networking component.

| **Cluster Index** | **Annotation** | **Putative Chemical Name** | **Inchi** | **SMILES** | **Molecular Formula** | **Theoretical Monoisotopic Mass** | **Theoretical Monoisotopic *m/z*, [M+H]+** | **Observed Monoisotopic *m/z*, [M+H]+ (rounded to third decimal)** | **Mass Error (ppm)** |
|---|---|---|---|---|---|---|---|---|---|
| 1907188 and 190718 | 1 | Clindamycin | InChI=1S/C18H33ClN2O5S/c1-5-6-10-7-11(21(3)8-10)17(25)20-12(9(2)19)16- | CCCC1CC(C(NC(C(Cl)C)C2C(O)C(O)C(O)C(SC)O2)=O)N(C)C1 | C18H33ClN2O5S | 424.1799 | 425.1871 | 425.188 and 425.187 | 8.1167 and -0.2351 |

| | | | | | | | | | |
|---|---|---|---|---|---|---|---|---|---|
| 2 | | | 14(23)13(22)15(24)18(26-16)27-4/h9-16,18,22-24H,5-8H2,1-4H3,(H,20,25) | | | | | | 9 |
| 3309544 | 2 | Clindamycin-O-glucuronide | not applicable | not applicable | C24H41ClN2O11S | 600.212 | 601.2192 | 601.219 | -0.33266 |
| 1865518 | 3 | N-desmethyl clindamycin | InChI=1S/C17H31ClN2O5S/c1-4-5-9-6-10(19-7-9)16(24)20-11(8(2)18)15-13(22)12(21)14(23)17(25-15)26-3/h8-15,17,19,21-23H,4-7H2,1-3H3,(H,20,24) | CCCC1CC(C(NC(C(Cl)C)C2C(O)C(O)C(O)C(SC)O2)=O)NC1 | C17H31ClN2O5S | 410.1642 | 411.1714 | 411.171 | -0.97283 |
| 6908035 | 4 | Clindamycin sulfoxide | InChI=1S/C18H33ClN2O6S/c1-5-6-10-7-11(21(3)8-10)17(25)20-12(9(2)19)16-14(23)13(22)15(24)18(27-16)28(4)26/h9-16,18,22-24H,5-8H2,1-4H3,(H,20,25) | CCCC1CC(C(NC(C(Cl)C)C2C(O)C(O)C(O)C(S(C)=O)O2)=O)N(C)C1 | C18H33ClN2O6S | 440.1748 | 441.182 | 441.183 | 2.26664 |
| 6907884 | 5 | Clindamycin N-oxide | InChI=1S/C18H33ClN2O6S/c1-5-6-10-7-11(21(3,26)8-10)17(25)20-12(9(2)19)16-14(23)13(22)15(24)18(27-16)28-4/h9-16,18,22-24H,5-8H2,1-4H3,(H,20,25) | CCCC1CC(C(NC(C(Cl)C)C2C(O)C(O)C(O)C(SC)O2)=O)[N+](C)([O-])C1 | C18H33ClN2O6S | 440.1748 | 441.182 | 441.183 | 2.26664 |
| 1951461 | 6 | N-desmethyl clindamycin sulfoxide | InChI=1S/C17H31ClN2O6S/c1-4-5-9-6-10(19-7-9)16(24)20-11(8(2)18)15-13(22)12(21)14(23)17(26-15)27(3)25/h8-15,17,19,21-23H,4-7H2,1-3H3,(H,20,24) | CCCC1CC(C(NC(C(Cl)C)C2C(O)C(O)C(O)C(S(C)=O)O2)=O)NC1 | C17H31ClN2O6S | 426.1591 | 427.1663 | 427.169 | 6.32072 |
| 2228486 | 7 | Clindamycin sulfone | InChI=1S/C18H33ClN2O7S/c1-5-6-10-7- | CCCC1CC(C(NC(C(Cl)C)C2C(O)C(O)C(O)C(S(C)(=O)=O)O2)=O)N(C)C1 | C18H33ClN2O7S | 456.1697 | 457.1769 | 457.177 | 0.21873 |

| | | | | | | | | | |
|---|---|---|---|---|---|---|---|---|---|
| | | | 11(21(3)8-10)17(25)20-12(9(2)19)16-14(23)13(22)15(24)18(28-16)29(4,26)27/h9-16,18,22-24H,5-8H2,1-4H3,(H,20,25) | | | | | | |
| 2228535 | 8 | Clindamycin sulfoxide N-oxide | InChI=1S/C18H33ClN2O7S/c1-5-6-10-7-11(21(3,26)8-10)17(25)20-12(9(2)19)16-14(23)13(22)15(24)18(28-16)29(4)27/h9-16,18,22-24H,5-8H2,1-4H3,(H,20,25) | CCCC1CC(C(NC(C(Cl)C)C2C(O)C(O)C(O)C(S(C)=O)O2)=O)[N+](C)([O-])C1 | C18H33ClN2O7S | 456.1697 | 457.1769 | 457.178 | 2.40607 |
| 6941163 | 9 | Putative metabolite 9 | not applicable | not applicable | C17H31ClN2O7S | 442.154 | 443.1612 | 443.162 | 1.80521 |

**Table S3.** R group table for putatively annotated clindamycin-related chemical **2**

| | $R_1$ | $R_2$ | $R_3$ |
|---|---|---|---|
| **2 -** Structure A | -glucuronide conjugate (C6H8O6) | -H | -H |
| **2 -** Structure B | -H | -glucuronide conjugate (C6H8O6) | -H |
| **2 -** Structure C | -H | -H | -glucuronide conjugate (C6H8O6) |

### Supplemental Figures

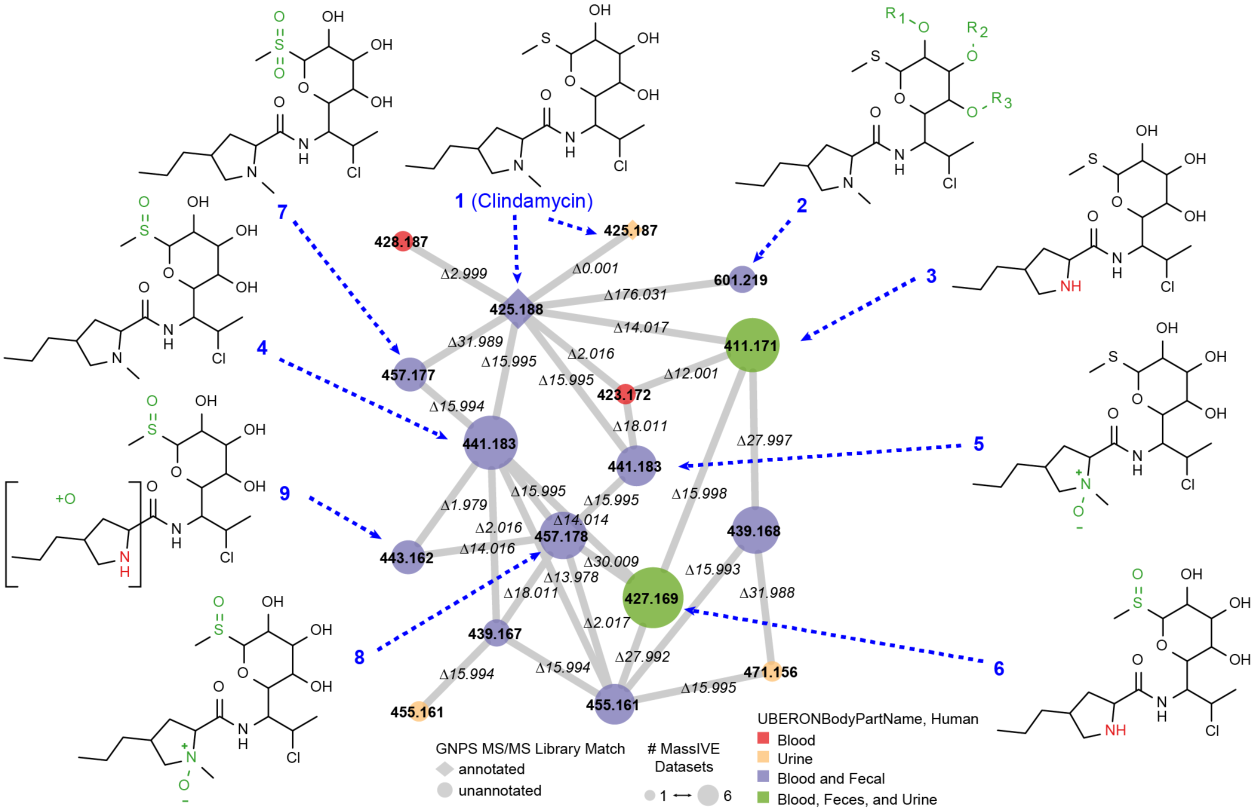


**Fig S1.** Molecular networking component associated with clindamycin (**1**) is displayed with putatively annotated clindamycin metabolites (**2-9**) observed in human blood, human fecal, and human urine data analyzed using GNPS' molecular networking. The location of modification on each putatively annotated clindamycin metabolites is indicated by color reflecting additions or subtractions from clindamycin's structure. The neutral charge structure for each chemical is displayed, whereas a positively charged version was detected in the molecular network. Further details on the annotated compounds are tabulated in **Table S2**. Note, the location of glucuronidation is ambiguous in **2** with possible locations indicated by $R_1$, $R_2$, and $R_3$ with corresponding information **Table S3**. The location of the oxygen atom addition in structure **9** could not be unambiguously determined, but exists within the structure in brackets.

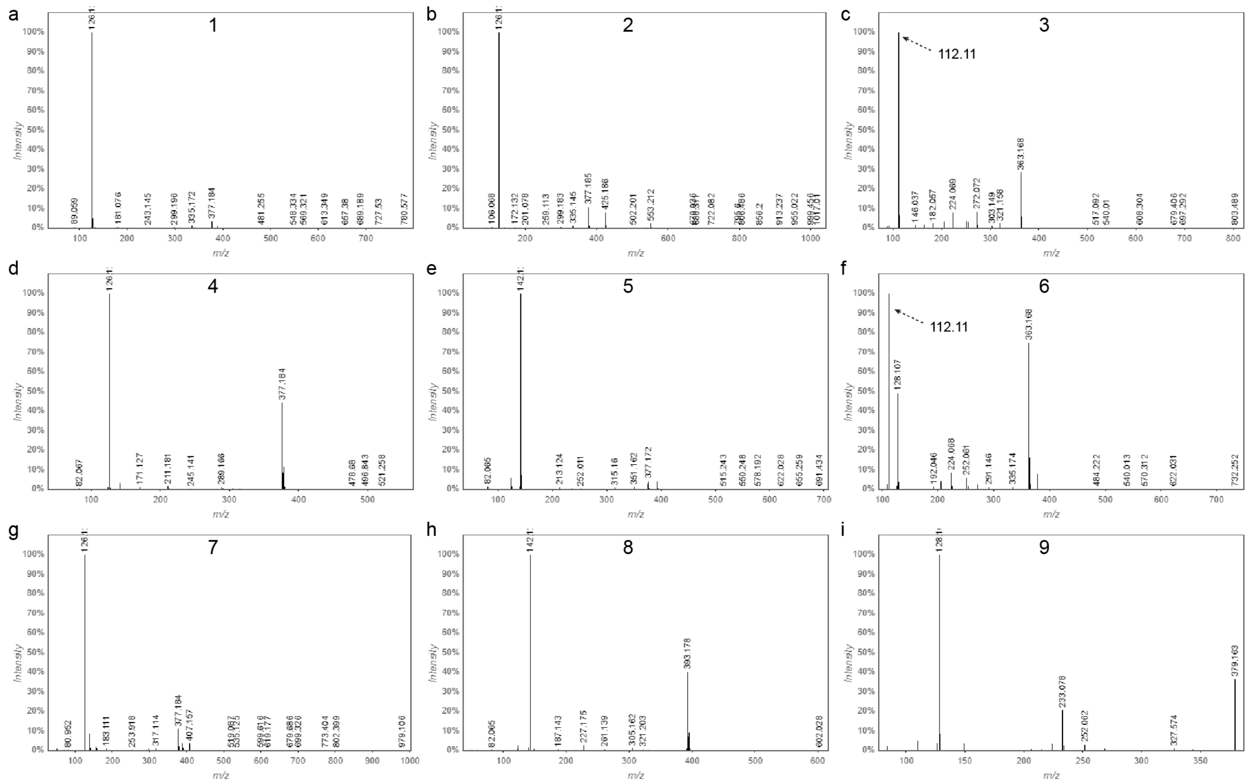


**Fig S2. (a)** MS/MS consensus spectrum annotated as clindamycin (**1**) based on spectral library matching. The MS/MS consensus spectrum for putatively annotated clindamycin metabolites **2-9** are displayed in panel (b) through (i), respectively. The product ion *m*/*z* 126 indicates that the substituted pyrrolidine in clindamycin's structure remains unmodified (such as in **1**, **2**, **4**, and **7**), the product ion *m*/*z* 112 indicates a subtraction of a methyl group from the substituted pyrrolidine (**3** and **6**), the product ion *m*/*z* 128 indicates the replacement of the N-methyl group with that of -OH in the substituted pyrrolidine (**9**), and the product ion *m/z* indicates the addition of an oxygen atom believed to originate from the formation of an N-oxide in the substituted pyrrolidine (**5** and **8**). The nitrogen rule was also applied and support the assignment of the substituted pyrrolidine product ions. The product ions corresponding to the neutral loss of $CH_4S$, $CH_4OS$, and $CH_4O_2S$ (from the sulfide, sulfoxide, and sulfone, respectively) produced the same product ion *m*/*z* 377 indicating the location of oxygen atoms. The glucuronide metabolite (**2**) was annotated based on a characteristic neutral loss corresponding to a C6H8O6 and product ions matching that of clindamycin. Further details on the annotated compounds are tabulated in **Table S2**.

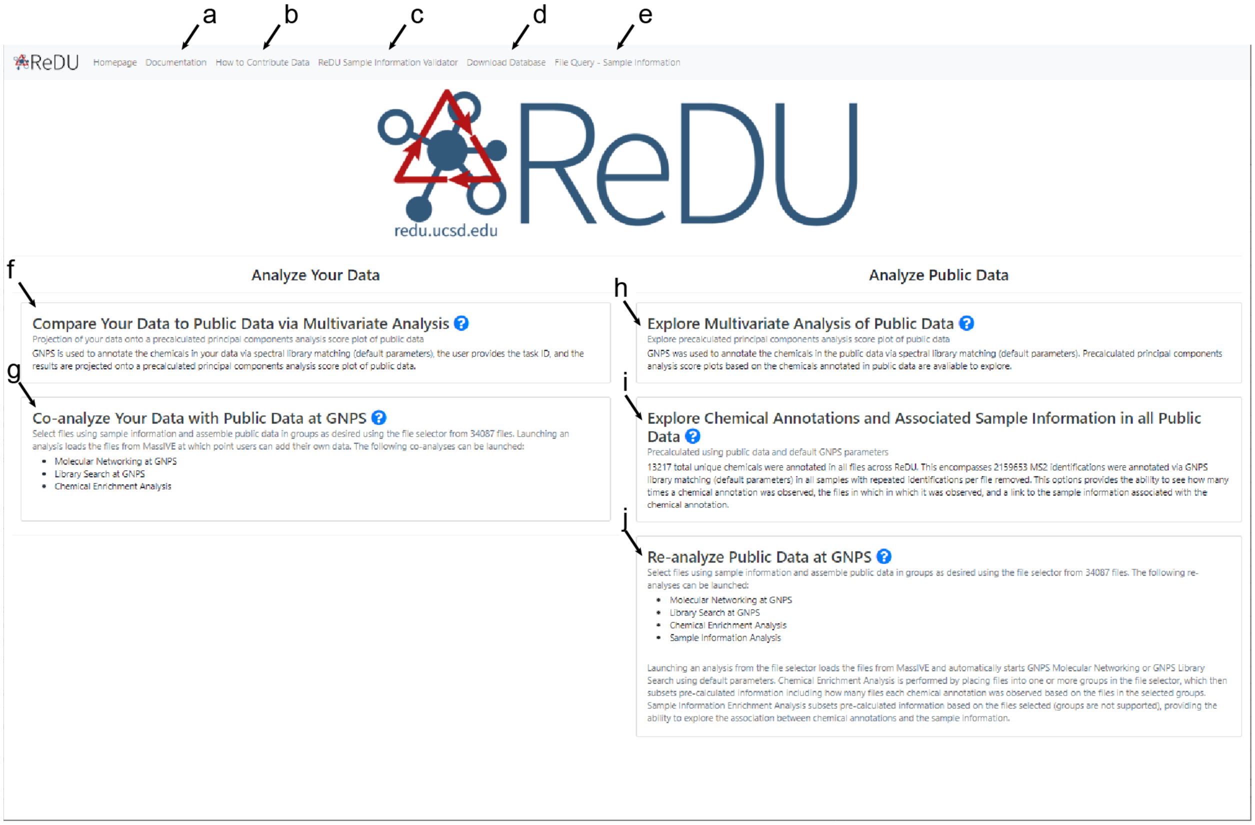


**Fig S3.** Screenshot of the ReDU website (redu.ucsd.edu) homepage with the following functionalities: (a) ReDU documentation, (b) information on data and sample information contribution, (c) sample information validator, (d) download ReDU sample information database, (e) Query the sample information in the ReDU sample information database given a filename in ReDU, (f) Project your data onto a precalculated PCA plot given a GNPS library search or molecular networking task ID, (g) Assemble cohorts of public files using sample information as well as include your data for molecular networking or library search, (h) Explore precalculated PCA plot generated from ReDU data, (i) Explore chemical annotation associations in ReDU data, and (j) Assemble cohorts of public files using sample information and launch molecular networking, library search, chemical enrichment analysis, or sample information association on the selected data.

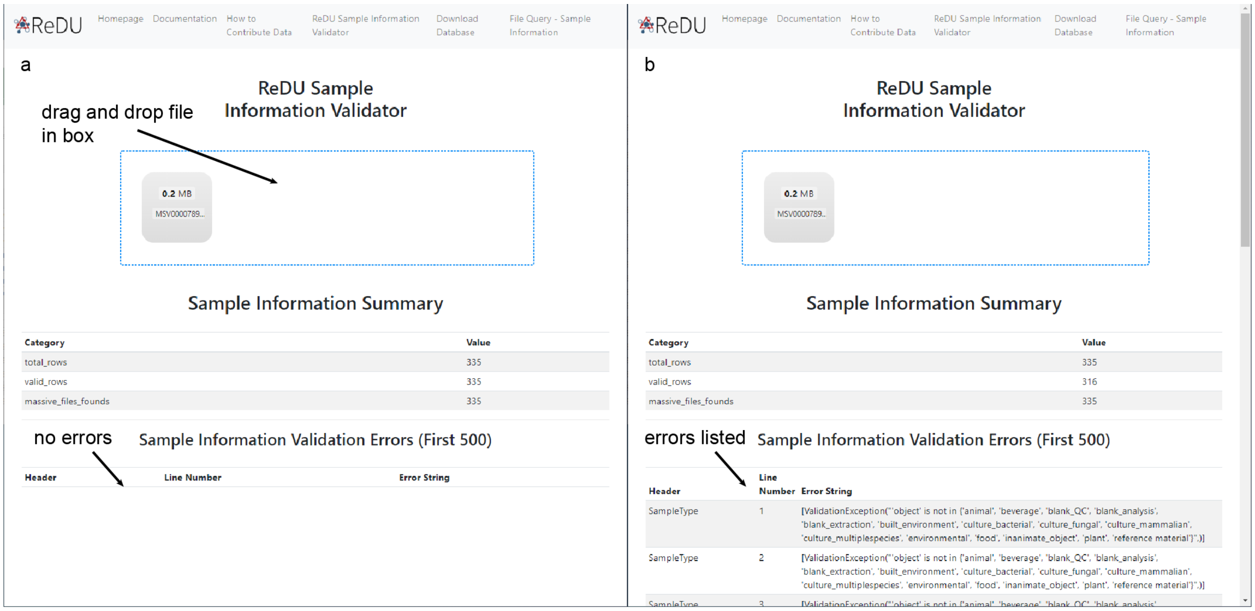


**Fig S4.** (a) Screenshot of validator when file passes. (b) Screenshot of validator when file fails.

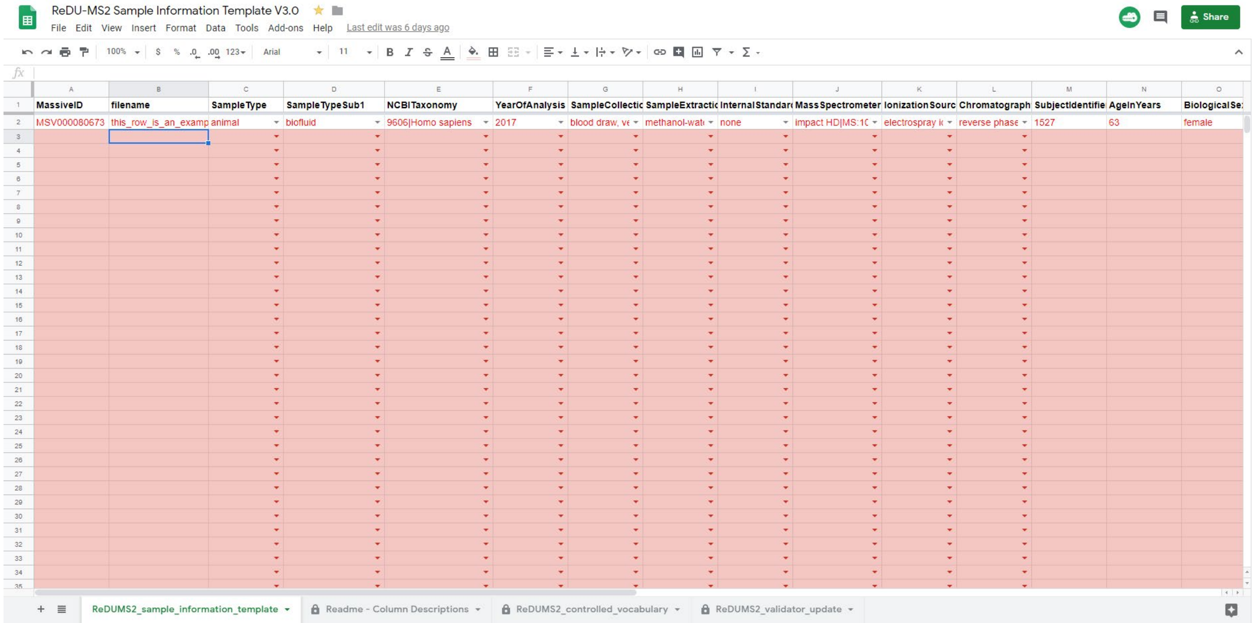


**Fig S5.** Screenshot of ReDU sample information template.

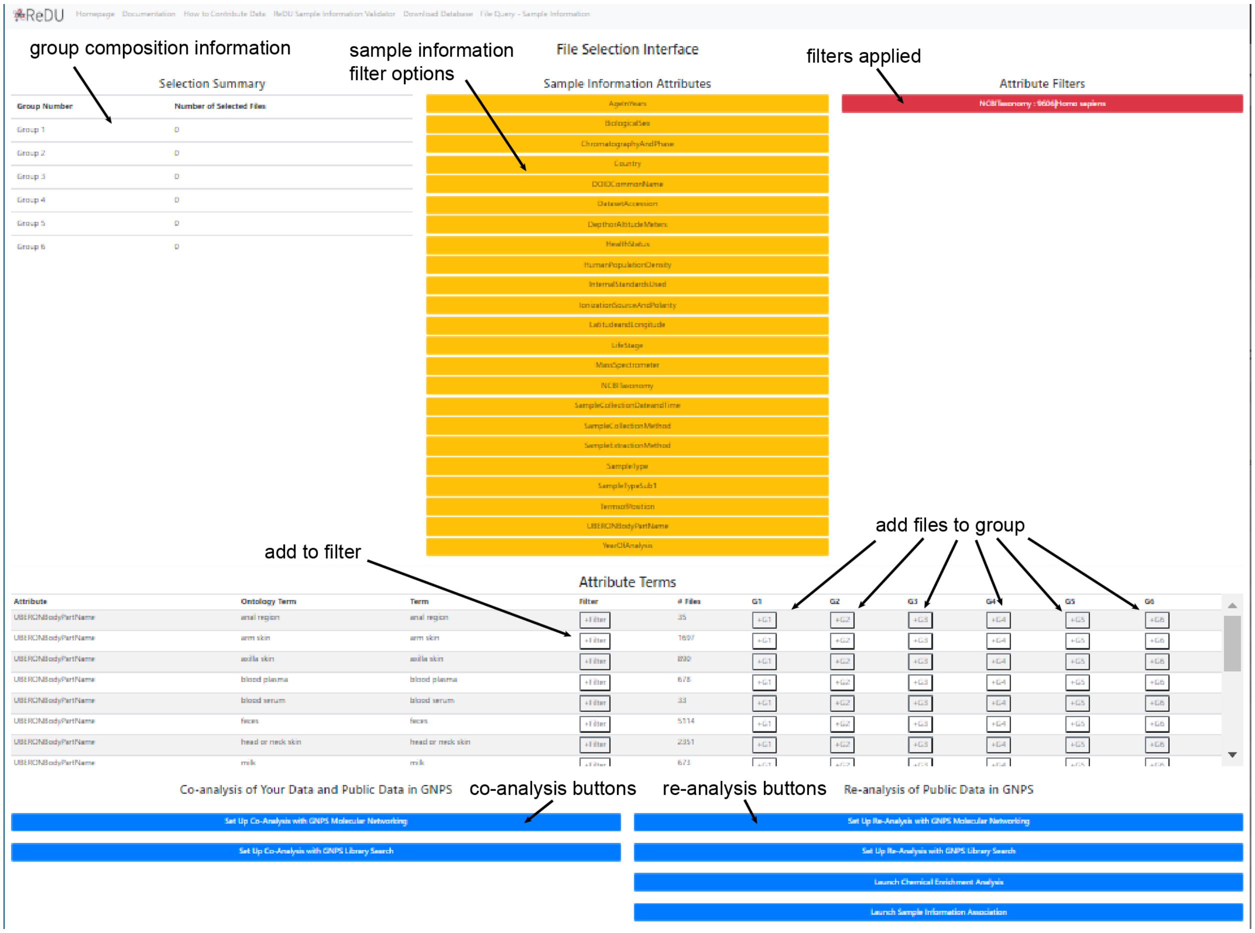


**Fig S6**. Screenshot of ReDU file selector.

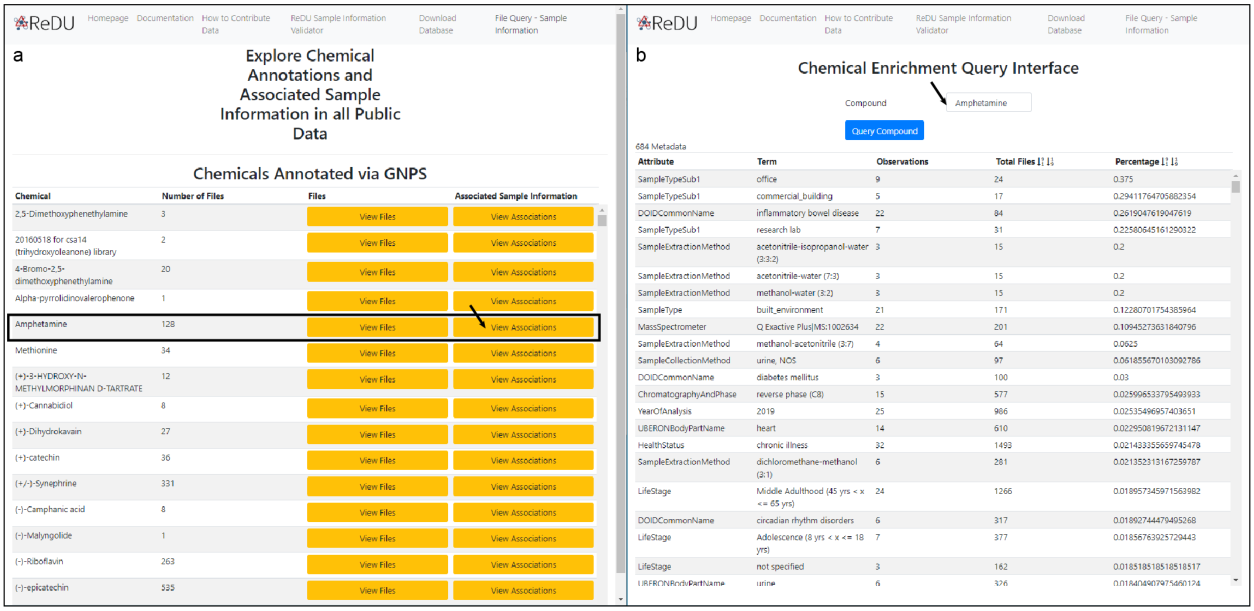


**Fig S7.** (a) Screenshot of the explore "Chemical Annotations and Associated Sample Information in all Public Data" page, **Fig S3i**. Users can search the table using Ctrl+F, illustrated by the box surrounding Amphetamine. Clicking on the "View Association" button directs you to (b) the screenshot of the sample information association table computed for that chemical (indicated in the field at the top of the page). Users can copy and paste this information for further processing and visualization.

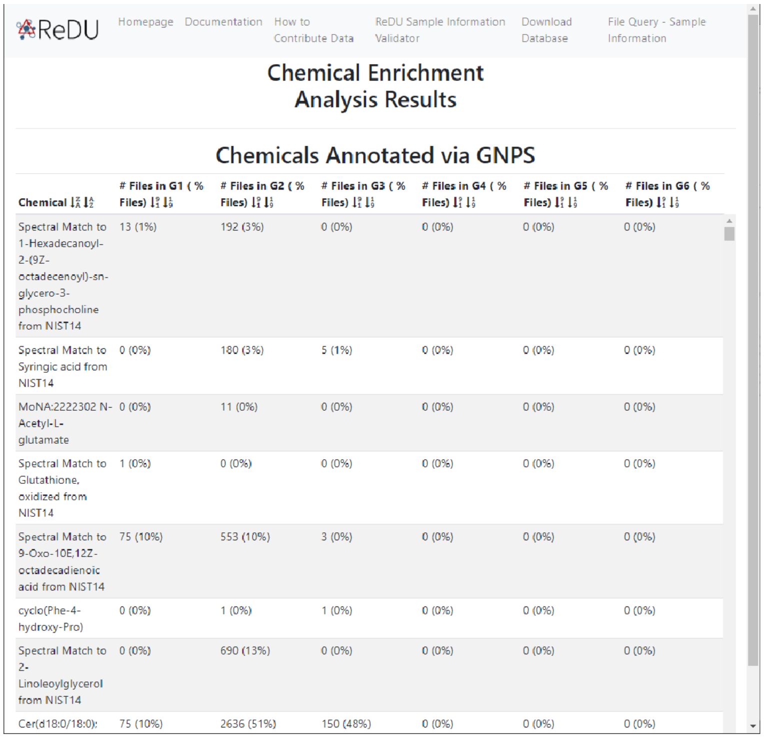
ReDU | Homepage | Documentation | How to Contribute Data | ReDU Sample Information Validator | Download Database | File Query - Sample Information

Chemical Enrichment Analysis Results

Chemicals Annotated via GNPS

| Chemical | # Files in G1 ( % Files) | # Files in G2 ( % Files) | # Files in G3 ( % Files) | # Files in G4 ( % Files) | # Files in G5 ( % Files) | # Files in G6 ( % Files) |
|---|---|---|---|---|---|---|
| Spectral Match to 1-Hexadecanoyl-2-(9Z-octadecenoyl)-sn-glycero-3-phosphocholine from NIST14 | 13 (1%) | 192 (3%) | 0 (0%) | 0 (0%) | 0 (0%) | 0 (0%) |
| Spectral Match to Syringic acid from NIST14 | 0 (0%) | 180 (3%) | 5 (1%) | 0 (0%) | 0 (0%) | 0 (0%) |
| MoNA:2222302 N-Acetyl-L-glutamate | 0 (0%) | 11 (0%) | 0 (0%) | 0 (0%) | 0 (0%) | 0 (0%) |
| Spectral Match to Glutathione, oxidized from NIST14 | 1 (0%) | 0 (0%) | 0 (0%) | 0 (0%) | 0 (0%) | 0 (0%) |
| Spectral Match to 9-Oxo-10E,12Z-octadecadienoic acid from NIST14 | 75 (10%) | 553 (10%) | 3 (0%) | 0 (0%) | 0 (0%) | 0 (0%) |
| cyclo(Phe-4-hydroxy-Pro) | 0 (0%) | 1 (0%) | 1 (0%) | 0 (0%) | 0 (0%) | 0 (0%) |
| Spectral Match to 2-Linoleoylglycerol from NIST14 | 0 (0%) | 690 (13%) | 0 (0%) | 0 (0%) | 0 (0%) | 0 (0%) |
| Cer(d18:0/18:0): | 75 (10%) | 2636 (51%) | 150 (48%) | 0 (0%) | 0 (0%) | 0 (0%) |

**Fig S8.** Screenshot of chemical enrichment results tabulated for human blood, feces, and urine samples in G1, G2, and G3, respectively.
